# Sharing individualised template MRI data for MEG source reconstruction: a solution for open data while keeping subject confidentiality

**DOI:** 10.1101/2021.11.18.469069

**Authors:** Mikkel C. Vinding, Robert Oostenveld

## Abstract

The increasing requirements for adoption of FAIR data management and sharing original research data from neuroimaging studies can be at odds with protecting the anonymity of the research participants due to the person-identifiable anatomical features in the data. We propose a solution to this dilemma for anatomical MRIs used in MEG source analysis. In MEG analysis, the channel-level data is reconstructed to the source-level using models derived from anatomical MRIs. Sharing data, therefore, requires sharing the anatomical MRI to replicate the analysis. The suggested solution is to replace the individual anatomical MRIs with individualised warped templates that can be used to carry out the MEG source analysis and that provide sufficient geometrical similarity to the original participants’ MRIs.

First, we demonstrate how the individualised template warping can be implemented with one of the leading open-source neuroimaging analysis toolboxes. Second, we compare results from four different MEG source reconstruction methods performed with an individualised warped template to those using the participant’s original MRI. While the source reconstruction results are not numerically identical, there is a high similarity between the results for single dipole fits, dynamic imaging of coherent sources beamforming, and atlas-based virtual channel beamforming. There is a moderate similarity between minimum-norm estimates, as anticipated due to this method being anatomically constrained and dependent on the exact morphological features of the cortical sheet.

We also compared the morphological features of the warped template to those of the original MRI. These showed a high similarity in grey matter volume and surface area, but a low similarity in the average cortical thickness and the mean folding index within cortical parcels.

Taken together, this demonstrates that the results obtained by MEG source reconstruction can be preserved with the warped templates, whereas the anatomical and morphological fingerprint is sufficiently altered to protect the anonymity of research participants. In cases where participants consent to sharing anatomical MRI data, it remains preferable to share the original defaced data with an appropriate data use agreement. In cases where participants did not consent to share their MRIs, the individualised warped MRI template offers a good compromise in sharing data for reuse while retaining anonymity for those research participants.

## 1 Introduction

There is an increasing demand for sharing data to improve reproducibility and efficiency in human neuroimaging studies according to FAIR practices (Findability, Accessibility, Interoperability, and Reusability). At the same time, researchers face ethical and legal responsibility and need to maintain the confidentiality of their research participants (White et al., 2020, Eke et al., 2021). The *Declaration of Helsinki* states that *“[e]very precaution must be taken to protect the privacy of research subjects and the confidentiality of their personal information”* (World Medical Association, 2013). The practice and principles of openly sharing scientific data conflicts with the need to protect the research participants. If we do not take adequate precautions when sharing neuroimaging data, we risk compromising the confidentiality of the research participants and losing their trust and willingness to contribute as volunteers.

The ethical conflict is in particular prevalent in neuroimaging. Human neuroimaging data, such as magnetic resonance imaging (MRI), magnetoencephalography (MEG), electroencephalography (EEG), positron emission tomography (PET), among others, does contain information that can potentially compromise the confidentiality of the research participant when shared unmodified and without constraints. Anatomical MRI data is the most identifiable, as it literally is an image of the research participant, including facial details that can be matched to photo databases (Schwarz et al., 2019). Task-related functional data from MEG, EEG, PET, and similar neuroimaging modalities are inherently less identifiable than MRI. It is, however, common in the analysis of MEG, EEG, and PET data to co-register the native functional data to the anatomical MRI. In EEG and MEG, this is used for source reconstruction, i.e., for mapping the measured channel-level activity onto the brain. Sharing the complete raw data from a study that uses any of these functional methods in combination with MRI, therefore, also needs to share the anatomical MRI.

In this paper, we focus on the sharing of neuroimaging data from MEG studies. MEG data itself consists of the time series of the magnetic fields or magnetic gradients measured with many sensors around the head. One of the strengths of MEG is that the magnetic fields originating from the synchronous postsynaptic potentials are virtually unaffected by the intermediate tissue between the brain and the sensors (Hari & Puce, 2017). This feature makes it possible to give a relatively precise estimate of the sources of the magnetic fields measured around the scalp. Thanks to the higher spatial resolution of MEG compared to EEG, but also since MEG channels are not placed on standardised positions relative to the participants head, it is common in MEG studies to use source reconstruction to localise the measured signals and experimental effects in the brain (Baillet, 2017). Source reconstruction requires a volume conduction model of the head that specifies how the electric and magnetic fields propagate in the brain and other tissues of the head. These models are created from structural MRIs of each research participant. Hence, MEG data is usually complemented with an anatomical MRI of each research participant. An independent researcher who wishes to reuse the shared MEG data to replicate or extend the original study will need the MRIs of the research participants. However, sharing the MRI data can potentially compromise the confidentiality of the research participants.

There are ways to approach a solution to the conflicting demand between open data and participant confidentiality. One solution is to withhold the directly identifiable data, such as the unmodified structural MRI. Previous recommendation for sharing neuroimaging data (predominantly fMRI) has been to share, e.g., statistical maps at the group level (Gorgolewski et al., 2015) or the peak locations of the activity (Fox & Lancaster, 2002). The processed data can be shared without a risk to compromise the confidentiality of the research participants. However, sharing already processed data limited the usability of the data beyond the replication of the study for which the data was collected. To (re)use data for testing new hypotheses, exploratory research, or testing out new analysis tools, one needs data that has not been processed. For usability of the data, it is most desirable to share as “raw” data as possible so that other researchers can replicate the entire processing pipeline or apply the preprocessing procedures most suited for their studies. The prevailing question is how “raw” data can be when sharing and still keep the participants anonymous in the data?

When sharing neuroimaging data, it is common to work with pseudonyms and remove all traces of the participants’ names and other directly identifiable information from the metadata. Furthermore, it is standard procedure to “deface” structural MRI before sharing the data, i.e., removing the parts of the image containing the face or skull-strip and removing all features except the brain (Theyers et al., 2021). However, though the face is easily the most identifiable feature, it is not the only feature of an MRI that can be used to identify individuals (Amico & Goñi, 2018; Byrge & Kennedy, 2019). The brain’s anatomy is unique to each person: the folding pattern of the cortex is identifiable, just like a person’s fingerprint, and could therefore be used to de-anonymise research participants if that brain fingerprint could be linked to another database (Aloui et al., 2018).

Sharing the anatomical MRIs for MEG source analysis is subject to the ethical consideration of sharing person sensitive information and the trade-off between replicability and participant confidentiality: on one end, share all data with the original MRI—on the other end, share only the MEG data and a template head model. Sharing the original MRI allows replication and to improve upon the entire MEG data analysis procedure and is, therefore, the most desirable from an open science perspective. However, this also makes the individual traceable anatomical information available, which may conflict with the informed consent that the participant provided. An intermediate solution is to share only the end-point of the MEG-specific anatomical MRI processing steps, which is the relatively coarse geometrical description of the volume conduction model. The MEG volume conduction model does not require facial details (Stenroos et al., 2014) but requires that all coregistration procedures are applied to the shared data, as, for example, done in the Human Connectome Project (Larson-Prior et al., 2013). This strategy is sufficient to reproduce certain types of MEG source analysis and is much less identifiable than the raw MRI, but can pose challenges if the re-analysis is done in other software such as MNE-Python (Gramfort et al., 2013), which expects to start from an anatomical MRI rather a co-registered headmodel.

Sharing only a template structural MRI, or data that has been transformed to template space, is another option for sharing functional neuroimaging data. The use of a template anatomical MRI is sometimes required in cases where individual MRIs cannot be acquired due to practical or ethical constraints, such as in children (Xu et al., 2019; Tadel, 2021). In this case, there is no person-sensitive individual anatomical information in the dataset. However, the replication of the analysis will depend on the accuracy or mismatch between the template and individual MRI, most notably when there are significant discrepancies in head size between the subject and the template. Considering a conventional SQUID-based MEG system, if the subject’s head is larger than the template, the sources will appear more superficial in the template than they are, or even outside the subject’s head. If the subject’s head is smaller than the template, the sources will appear deeper in the template than they are. If source localisation is of interest to the MEG study, the size mismatch therefore needs to be considered. Possible solutions in this case are to scale the MRI template to match head surface points that are recorded using a 3D tracker, such as a Polhemus, or to scale the template to match the individual participant’s head circumference.

In the following, we outline a pipeline for processing anatomical MRI data for MEG source analysis that aims to strike a compromise between anonymising the research participants and maximising the value of the shared data for researchers that want to reuse it. The pipeline is based on warping a template anatomical MRI to the individual anatomical MRI of the research participants. The approach consists of nonlinear deformation of the template MRI to align with the subject’s MRI. Such nonlinear transformation procedures are widely available and commonly used to analyse source reconstructed MEG data, but in the opposite direction: they are frequently used to warp from the subject’s native MRI space to a template or reference brain (e.g., Talairach or MNI) for group-level inference. The procedure we describe results in a warped MRI template that is adequate for creating volume conductor models for MEG source analysis but does not provide precise anatomical information about the subject’s cortical folding patterns. It can readily be implemented with available analysis software—it is only a matter of adding the extra step of warping a template MRI as outlined in Figure 2. The warped templates effectively remove the individually traceable information: they are, after all, not images of the participant.

Although the head model—and thereby the source reconstructions—will not be numerically identical to those based on the original MRI data, we hypothesise that the results of re-analysis are still within a range that complies with the principles of replicability. Therefore, we expect this procedure to offer a compromise between FAIR data sharing and keeping the confidentiality of the research participants, especially for MEG researchers that currently are not allowed to share data due to ethical or legal constraints or hesitations of their institutes. The procedure is foremost aimed at constructing the volume conduction model and source reconstruction methods that merely depend on the 3-D volumetric information about the source compartment (e.g., dipole analysis, beamformers), and we expect it to be less adequate for methods that require detailed cortical sheets (e.g., distributed source reconstruction methods such as minimum-norm estimates).

In this study, we present in detail the procedure for creating individual subject warped template MRIs that can be shared while providing greater anonymity for the research participants. Furthermore, we evaluate the performance by comparing the similarity between source reconstruction using the original MRI and a warped template with four types of MEG source reconstruction. The source reconstruction methods represent typical methods used in MEG research and are presented in ascending order of complexity in terms of model constraints and dependence on anatomical features from the MRI. First, we compared single equivalent current dipole fits as the geometrically least constrained model that we expect to be the least affected. Second, we compared reconstructions using the dynamic imaging of coherent sources (DICS) beamformer method (Gross et al., 2001), using a full-brain regular 3-D grid for the sources. Third, we compared virtual channels estimated with an LCMV beamformer (Van Veen et al., 1997), using an atlas to define cortical parcels as the regions of interest. Finally, we tested the similarity between minimum-norm estimates (MNE) of distributed source models constrained by the location on the cortical sheets (Hämäläinen & Ilmoniemi, 1994). In this case, the cortical sheets are obtained from either the original MRI or the warped template. Since the MNE depends on the anatomical information, we expect the anatomical discrepancies between the original MRI and the warped template to cause the most considerable difference in source estimation for this approach. Considering all four source reconstruction methods: if the discrepancy between source reconstruction with a warped template and the subject’s actual MRI is sufficiently small, we argue that this procedure provides a useful compromise between the need for sharing realistic MRIs and protecting the confidentiality of the research participants.

## 2 Methods

### 2.1 Data used in this study

The data used in the study is a MEG tutorial dataset obtained from a male participant (age 28) containing functional MEG recordings and a structural MRI. The data was acquired according to local ethical regulations at the Karolinska Institutet and is used for this study with the participant’s consent. The data is available at: https://zenodo.org/record/5053234.

The participants original MRI is a 3D T1-weighted magnetisation-prepared rapid gradient-echo (MPRAGE) sequence structural image (voxel size: 1 x 1 x 1 mm) obtained on a GE Discovery 3.0 T MR scanner. We used the *Colin27* template MRI (Holmes et al., 1998) as the basis to create the individualised templates.

MEG was recorded with a Neuromag Triux MEG scanner with 102 magnetometers and 204 planar gradiometers at a sample rate of 1000 Hz. The participant received 160 tactile stimulations to the index finger of the right hand at a rate of 0.3 Hz while watching a silent movie. Continuous HPI was measured for the duration of the recording.

The MEG was processed offline by first applying temporal signal space separation (tSSS) to suppress artefacts outside the scanner helmet and to correct for head movement during the recordings (Taulu & Simola, 2006). The tSSS had a buffer length of 10 s and a cut-off correlation coefficient of 0.98.

Offline MEG data processing was done with the FieldTrip toolbox (Oostenveld et al., 2011) in MATLAB (R2016b). The data was low-pass filtered at 100 Hz and cut into epochs from −2.0 to 2.0 seconds relative to the tactile stimulation and baseline corrected by the mean of the epoch. Bad trials were semi-manually rejected based on trial-by-trial variance. A total of 158 trials remained after data cleaning for the subsequent source analyses. The data was cut into epochs from −0.2 to 0.8 s relative to stimulation and baseline corrected with a baseline from −0.2 to 0.0 s before averaging trials to calculate the evoked responses (Figure 1).

**Figure 1:**
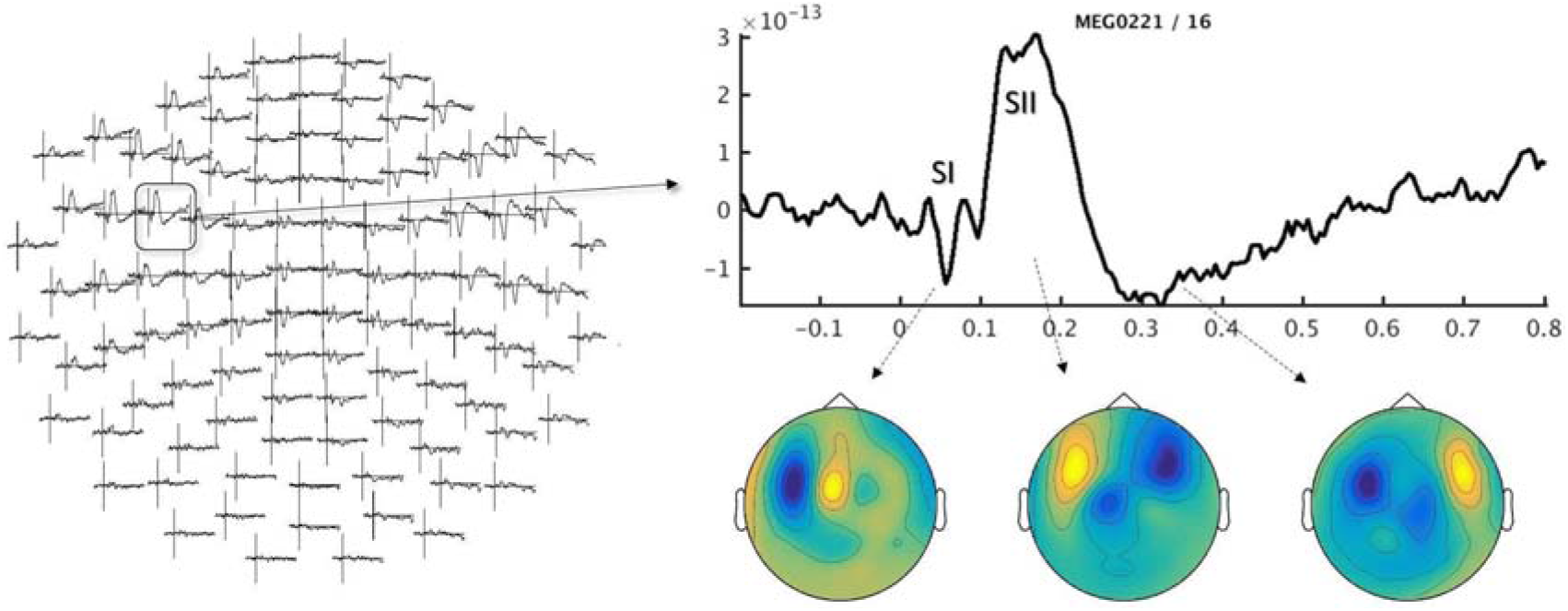
Sensor-level ERF to tactile stimulation in the MEG data used to compare source reconstruction using the original MRI and warped template, for the magnetometer sensor array (left), a representative sensor (right, top) and magnetometer topographies for the SI, SII, and late complex components in the response.

### 2.2 Creating individualised warped templates

The procedure for replacing the subject’s structural MRI with a warped temple is introduced as a preliminary step in the typical processing of MRI for MEG source reconstruction (Figure 2). It consists of a template MRI that is warped non-linearly to the subject’s MRI. The warped MRI can then be used in the following steps, as usual. Any neuroimaging analysis tool capable of warping MRI from one image to another can, in principle, be used for this procedure. In this demonstration and the accompanying tutorial scripts, the procedure was done with functions from SPM12 (www.fil.ion.ucl.ac.uk/spm/software/spm12) through the FieldTrip toolbox for MEG/EEG analysis.

**Figure 2:**
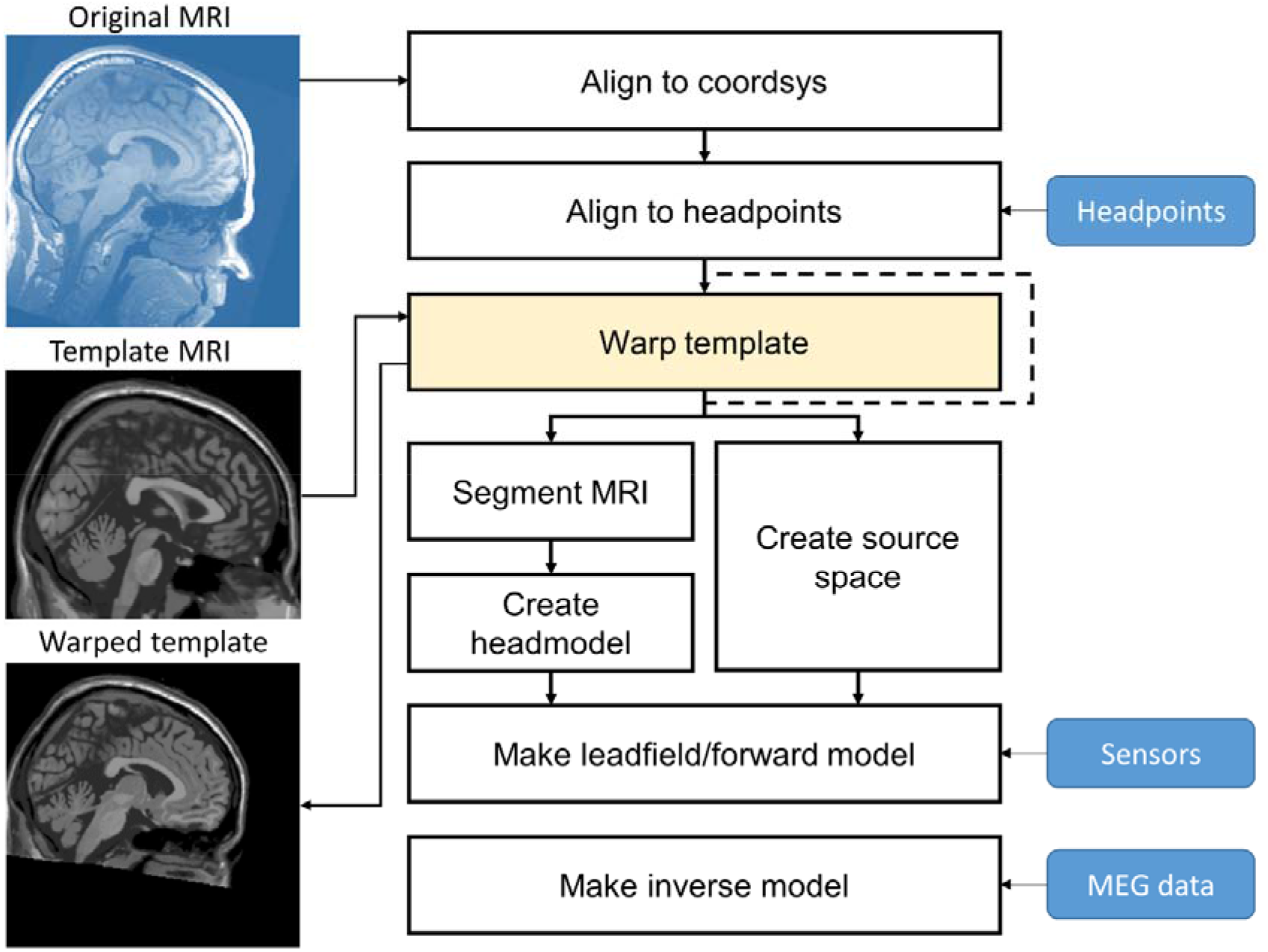
Overview of the procedure for creating individualised warped templates and where this step would enter the usual MRI processing pipeline in MEG source reconstruction. Displayed in blue is the actual data from the participant. The original MRI is imported in the first step and aligned to the MEG coordinate system. A template is then loaded in the second step and warped to the original MRI (yellow box). The warped template is used in the following processing and MEG source reconstruction in the exact same way as the original MRI. The yellow box is the additional step in the pipeline where the warping is done, and the dashed line indicates the pipeline using only the original MRI.

The subject’s MRI is realigned to the desired MEG system coordinate system (typical coordinate systems in MEG analysis are “Neuromag”, “CTF” and “ACPC” coordinate systems), then aligned to the head shape measured during the MEG recording, and subsequently resliced. The realigned volume is written to disk as the reference MRI that we will warp to in the next step.

In the second step, the template is warped to the subject’s MRI. The warping was done with the nonlinear spatial normalisation procedure in SPM12 (Ashburner & Friston, 1999). Warping a subject’s MRI to a standard template is a standard procedure in MRI normalisation. Here, we use the procedure to do the opposite and warped the *Colin27* template to the realigned original MRI. To facilitate the reverse normalisation of the template to the individual anatomy, we first segmented the subject’s MRI to create tissue probability maps (Ashburner & Friston, 2009) and used the individual tissue probability map for the nonlinear spatial normalisation.

The subsequent processing of the warped template was identical to that of the original MRI and followed a standard preprocessing pipeline of structural MRI for MEG source analysis from this stage and onwards. If the warped template provides a sufficient trade-off between anonymising data for data sharing and reproducibility of MEG source reconstruction results as proposed, we expect: 1) that the anatomical features of the warped template are sufficiently dissimilar from the original MRI, and 2) that the results from MEG source reconstruction using the warped template are sufficiently similar to the MEG source reconstructions using the original MRI.

To test whether MEG course reconstruction using the warped template would yield reliable results comparable to the same analysis using the original MRI, we processed the warped template and the original MRI with identical pipelines to compare the performance for different MEG source reconstruction methods.

The warped template and the original MRI were first aligned to the MEG coordinate system using points measured on the subject’s head during MEG acquisition with a Polhemus Fastrak 3D motion tracker. The volumes were then segmented into a single compartment containing the brain used to create single-shell volume conductor models (Nolte, 2003) for the original MRI and the warped template (see Figure 3).

**Figure 3:**
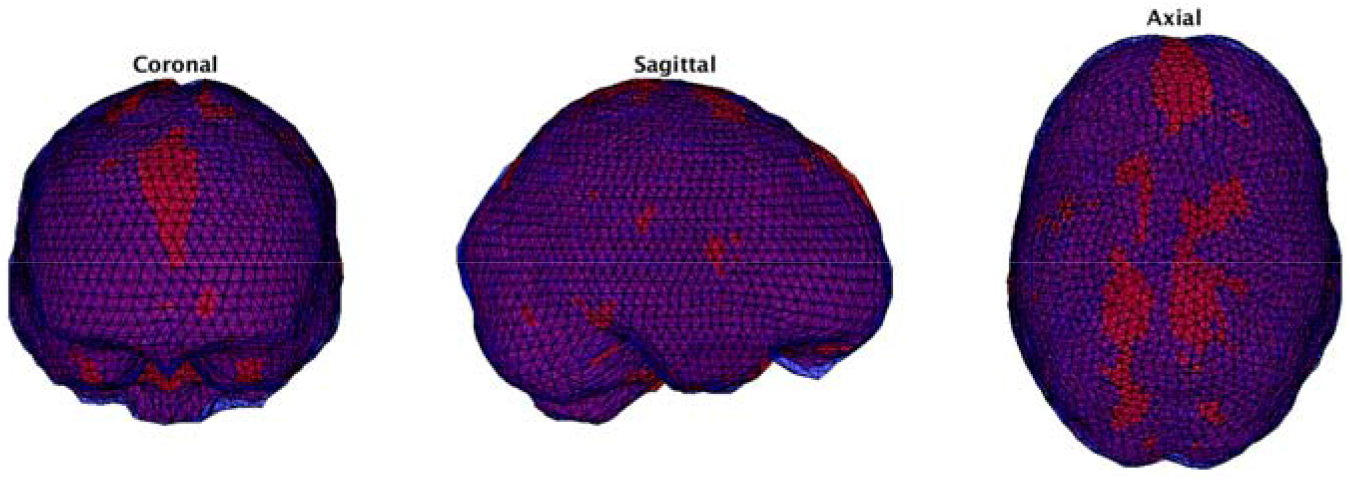
Volumetric head models created from the brain masks of the warped template (red) and the original MRI (blue) plotted within the same coordinate system.

### 2.3 Evaluating individualised templates

#### 2.3.1 Dipole analysis

The first type of MEG source reconstruction we evaluated were single equivalent current dipole models. First, we fitted a single dipole to the SI component (50-60 ms; see Figure 1) to test the localization agreement between the volume conductor models from the original MRI and the warped template. We estimated the dipole models separately for magnetometers and gradiometers. Following the same procedure, we fitted dual-dipole models to the SII component (115-155 ms; Figure 1).

Finally, we fixed a single equivalent current dipole to the SI dipole position and estimated the dipole moment for the whole time window from 0-500 ms relative to simulation; this allowed us to compare the estimated dipole moment time-series between two different volume conductor models.

#### 2.3.2 DICS beamformer

A strength of MEG is its ability to measure oscillatory brain responses and reconstruct its sources. We compared the reconstruction of oscillatory brain responses using the dynamic imaging of coherent sources (DICS) beamformer (Gross et al., 2001) to estimate the sources of the event-related beta desynchronization to the sensory stimulation. We selected a time window of interest from 220 ms to 500 ms and a baseline time window from −500 ms to −220 ms relative to the tactile stimulation. We first estimated the power spectral densities (PSD) and cross-spectral densities (CSD) between all pairs of sensors at 20 Hz using FFT after applying a single Hanning taper. The combined CSD of the time window of interest and of the baseline was used to create the DICS spatial filters for the original MRI head model and warped template.

The source spaces consisted of an evenly spaced 6×6×6 mm grid covering the entire volume defined in standard MNI coordinates. The MNI grid was warped to the original MRI and to the warped template and used for DICS source estimates for the two pipelines. The beamformer filters were applied to the CSD from the beta desynchronization and baseline time windows, respectively, and the source power in the beta desynchronization windows was normalised by the source power from the baseline time window. The normalised source estimates were then compared between the two head models.

#### 2.3.3 Atlas-based beamformer virtual channels

For the third comparison, we estimated “virtual channel” time series based on beamformer projection to pre-defined anatomical regions of interest (ROIs). The ROIs were defined from the automatic anatomical labelling (AAL) atlas (Tzourio-Mazoyer et al., 2002). The ROIs were the postcentral gyrus and precentral gyrus for both left and right hemispheres, representing the cortical areas where the SI and SII responses usually have peak values. In addition, we included an ROI in the inferior frontal lobe to represent a cortical area that is generally not associated with the processing of tactile stimuli. In addition to the cortical ROIs, we added a virtual channel in the left and right Areas 4/5 of the cerebellum. Though it is debated how much of the neural signals MEG picks up from the cerebellum, there is converging evidence that MEG can measure cerebellar signals at a lower signal-to-noise ratio than cerebral signals (Andersen & Lundqvist, 2019; Samuelsson et al., 2020). We, therefore, added cerebellar virtual channels to compare the similarity of signals reconstructed in this area using either the original MRI or the warped template. Finally, we added virtual channels in the left and right thalamus as a source in the centre of the brain that would have virtually no contribution to the measured MEG signal, and any reconstructed signal would be an expression of noise projection. The question that we address by the inclusion of the control ROIs, in which we expect no activity, is that even if the reconstructed signals do not represent actual neural signals, will the estimates nonetheless be the same for the two methods?

The anatomical atlas was projected onto a standard grid in MNI coordinates and then warped to the original MRI and to the warped template. The warped grids were used as source models to create beamformer filters using the LCMV beamformer (Van Veen et al., 1997) on the evoked data. The filters were then applied to the single-trial MEG data for the source locations defined in the ROI. Virtual channels were calculated for each ROI as the eigenvector with the largest eigenvalue of the time series of the source point within the ROI.

#### 2.3.4 Minimum Norm Estimates

For the minimum-norm estimate source reconstruction, we first exported the original MRI and warped template to Freesurfer (Fischl & Dale, 2000) to reconstruct the cortical sheet of the grey-white matter boundary. Source models were created by sampling 8004 points uniformly across the cortical surfaces of the original MRI and warped template. The points were sampled from the two surfaces by taking the points with the same indices in the Freesurfer template that the MRIs are aligned to during the surface extracting procedure in Freesurfer. This means that the points used in the source models correspond to the same standardised anatomical location, even if the geometrical locations are different between the warped template and the original MRI. This allows a direct comparison between the reconstructed activities on the downsampled surfaces.

We then applied source reconstruction to the evoked responses of the pre-whitened gradiometer evoked responses using noise weighted minimum-norm estimates (Dale et al., 2000). The noise covariance matrix was estimated from the baseline period of the epochs.

#### 2.3.5 Anatomical MRI features

To assess the dissimilarity between the anatomical MRIs, we performed a morphological analysis of the original MRI, the warped template, and the unmodified template. There will be a degree of both similarity and dissimilarity in the morphological features between the warped template and the original MRI. To what degree differences are sufficient to provide anonymity or what the degree of similarity allows to re-identify the research participant is difficult to quantify, as it depends on the external database that would be used for linkage attacks (e.g., the exact same MRI scan, another MRI scan of the same subject, or facial photos from a public database), the size of the sample available in the external database, the algorithm applied by those performing the re-identification attempts, and the context in which the re-identification attempt is made, i.e. what other data besides the warped template is available to those attempting the re-identification? Therefore, it is not possible to give a universal score for the “anonymity” of the warped templates; we only know for sure that the morphological features of the warped templates should be dissimilar to those of the original MRI. However, as we still expect some correlation between morphological features, we rationalised that the similarity between the warped template and the original MRI should not be greater than the similarity between the warped template and the original template.

To compare the morphological similarity between the warped template, the original MRI and the original template were processed with Freesurfer (Dale et al., 1999) to get summary statistics of the surface area, grey matter volume, average cortical thickness, and mean curvature within cortical parcels based on the anatomical labels provided by Freesurfer automatic labelling (Destrieux et al., 2010).

### 2.4 Comparing the results from the original MRI and warped template

To compare how similar the source reconstructions based on the warped template were to the source reconstructions based on the original MRI, we used Krippendorff’s α (Krippendorff, 1970) to quantify the reliability between the source estimates. In short, α is a generalised measure of the ratio between the observed disagreements between two (or more) measurements and the expected disagreement between the measurements at random given the observed values, which can be parametric, ordinal, or nominal data. A value of α = 1 means perfect agreement (i.e., all values are equal), whereas α = 0 means no agreement between measurements. There is no standard cut-off for what is considered an acceptable agreement as it depends on what would be deemed acceptable given the type of data. Based on recommendations for reliability analysis in the literature (Beckler et al., 2018; Koo & Li, 2016; Krippendorff, 2018), we use the following indicators as qualitative interpretation of α: α > 0.90 indicates excellent reliability, 0.80 < α < 0.90 indicate good reliability, 0.66 < α < 0.80 indicate moderate reliability, and α < 0.66 are considered poor reliability.

Krippendorff’s α for the dipole moment time series was calculated as the agreement across the whole time series by pairwise mapping the activity at each time point from 0-500 ms. For the DICS source reconstruction, Krippendorff’s α was calculated across the entire source space by pairwise mapping the estimated source power in standard MNI space. Krippendorff’s α for the virtual channels was calculated by pairwise mapping time points across the time series for each virtual channel. Krippendorff’s α for the MNE source reconstruction was calculated across the entire source space for all time points by matching the corresponding surface points and time points.

For the morphological features, Krippendorff’s α was calculated across the summary values from all parcels for respectively the surface area, grey matter volume, average cortical thickness, and mean curvature.

## 3 Results

### 3.1 Dipole analysis

The estimated dipole locations for the single dipole models of the SI component using the warped template and the original MRI had a difference of 0.6 mm for the dipole fits using magnetometers and 0.1 mm for the dipole fits using gradiometers. The difference in the estimated dipole locations for the symmetrical dual dipole models fitted to the SII component was 0.5 mm for the magnetometer fit and 0.2 mm for the gradiometer fit.

The dipole moment estimates from 0-500 ms and residual variance of the dipole models using fixed dipole location is shown in Figure 4. The agreement in the estimated dipole time series was very high—virtually identical as seen in Figure 4—with Krippendorff’s α = 0.999 for the magnetometer dipole time-series and α = 0.999 for the gradiometer dipole time-series.

**Figure 4:**
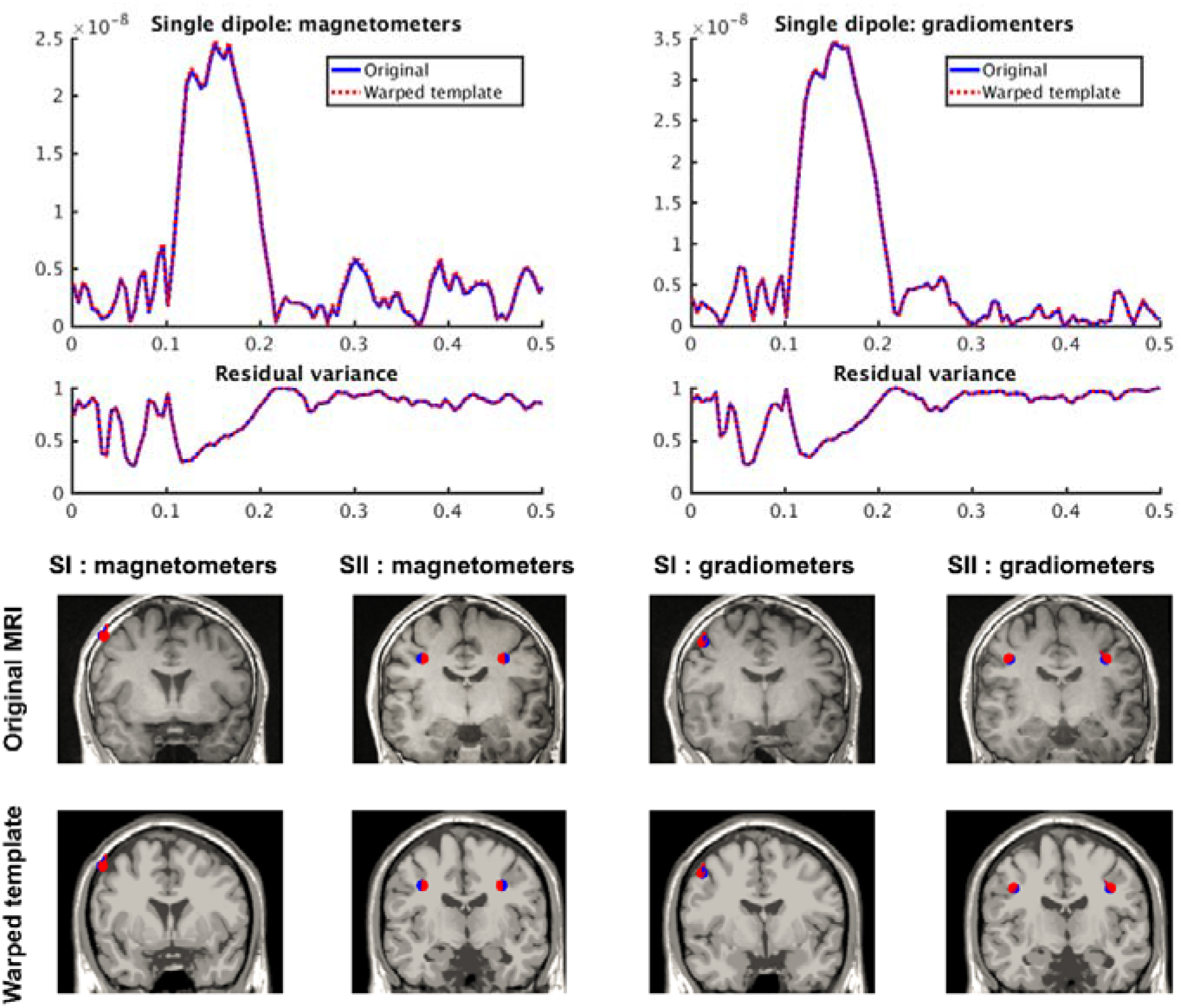
Source estimates from dipole fits. Top: Dipole power and residual variance of the dipole models from 0-500 ms relative to stimulation for single dipole fits to magnetometers (left) and gradiometers (right) using the original MRI (solid blue lines) and the warped template (dashed red lines). Bottom: dipole localisation for the dipole fits of the evoked SI and SII components with magnetometers (left) and gradiometers (right) for the original MRI (blue) and the warped template (red)

### 3.2 DICS beamformer

The whole-brain source reconstructions of the moment-related beta desynchronization using DICS (Figure 5) showed high similarity in the estimated source power for the original MRI and warped template (α = 0.995). The difference in source power between the estimates using the original MRI and the warped template did not show any systematic patches of difference.

**Figure 5:**
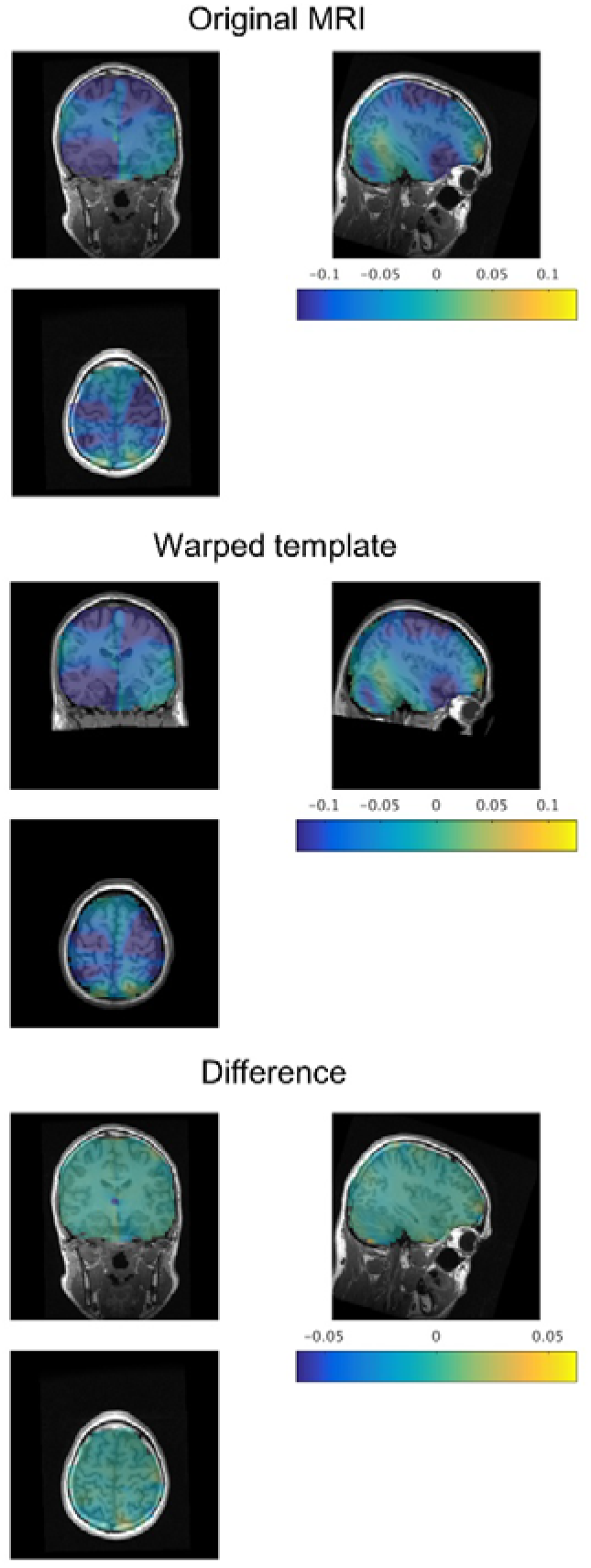
Topographic distribution of DICS source power for the movement-related beta desynchronization normalised by the baseline power. Top: using the original MRI. Middle: using the warped template. Bottom: the difference in the source estimates using the original MRI and warped template projected onto the original MRI.

### 3.3 Atlas-based beamformer virtual channels

The time series of the virtual channels estimated from the atlas-based beamformer showed overall high degrees of similarity for all ROIs, as shown in Figure 6. Krippendorff’s α was very high for all sensorimotor cortical ROIs that we expected to be the primary contributing sources to the evoked response (left precentral gyrus: α = 0. 994, right precentral gyrus: α = 0.998, left postcentral gyrus: α = 0.999, right postcentral gyrus: α = 0.997). Although the time series of the inferior frontal cortical virtual channels showed little activity as expected, it still showed excellent similarity (left: α = 0.998, right: α = 0.996). The cerebellar virtual channels also showed excellent similarity for both the left and right ROI (left: α = 0.995, right: α = 0.987). The similarity of the left thalamic virtual channel was comparable to the cortical sources (α = 0.993), but the source estimates in the right thalamic virtual channel showed a high systematic disagreement (α = −0.993), as seen in Fig 6. There was a systematic sign-flip in the polarity of the estimated signal, which can be explained by the sign of the eigenvector being arbitrary.

**Figure 6:**
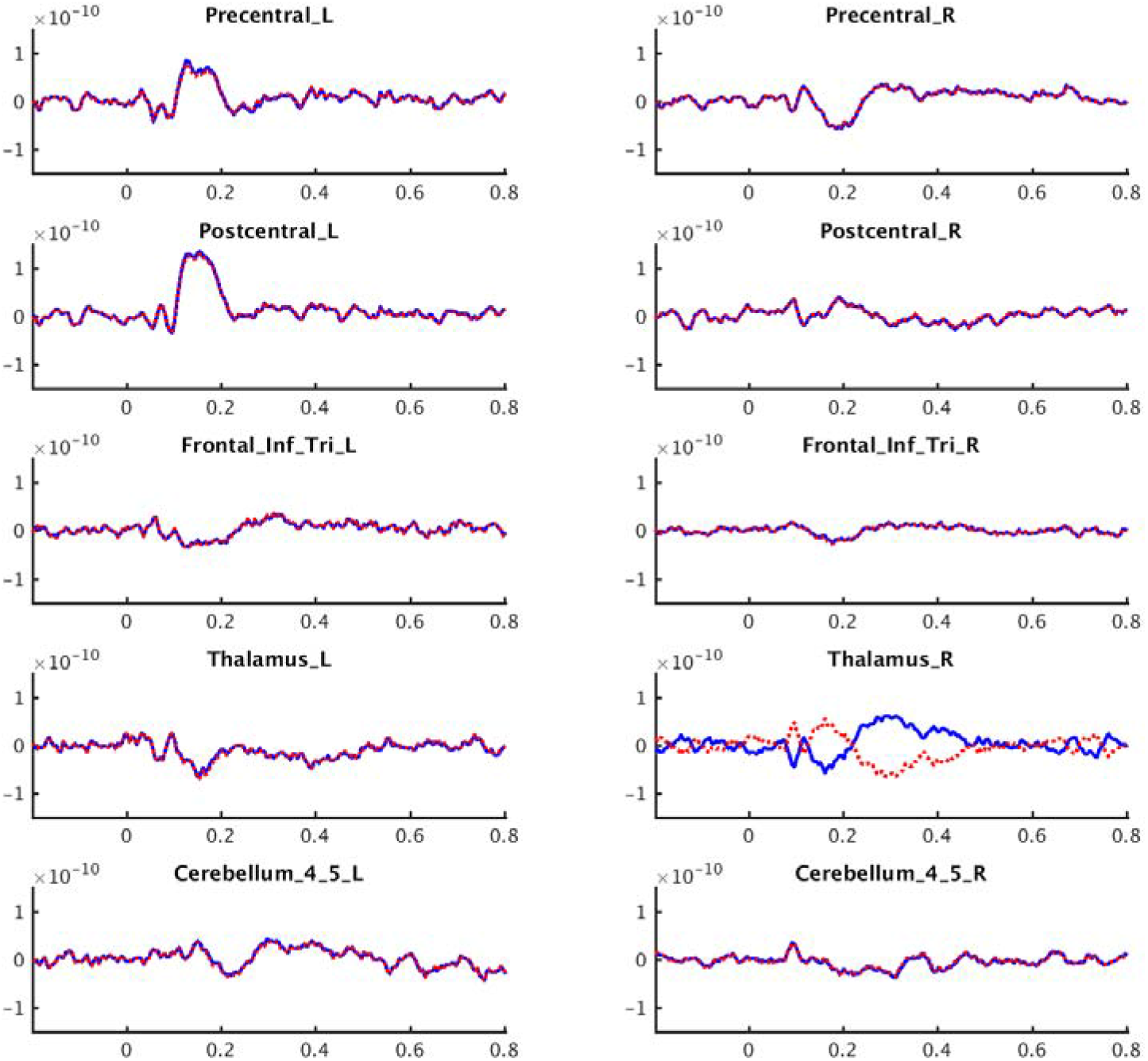
Virtual channel evoked responses at the ROIs using the warped template (red) and the original MRI (blue).

Overall, there was an excellent similarity between the estimated virtual channels with minimal numerical error. Even in virtual channels where we expected the estimated signals to reflect mostly noise, the two procedures produced numerically similar results, despite the sign-flip in one virtual channel.

### 3.4 Minimum Norm Estimates

The overall topography of the MNE source reconstruction was qualitatively alike. The evoked response to the tactile stimulation in both source reconstructions showed the maximum source activity in the sensorimotor cortex (Figure 7). However, the quantitative similarity analysis yielded only moderate agreement (α = 0.761), confirming that the difference in the cortical structure between the original MRI and the warped template compromises the source reconstruction’s numerical precision. Since the MNE source reconstruction estimates all source points to have non-zero power, any point that is estimated as arbitrarily close to zero in one case but slightly larger in the other case will therefore contribute to the discrepancy, even if both values would fall below what one would consider meaningful activation in the source estimate.

**Figure 7:**
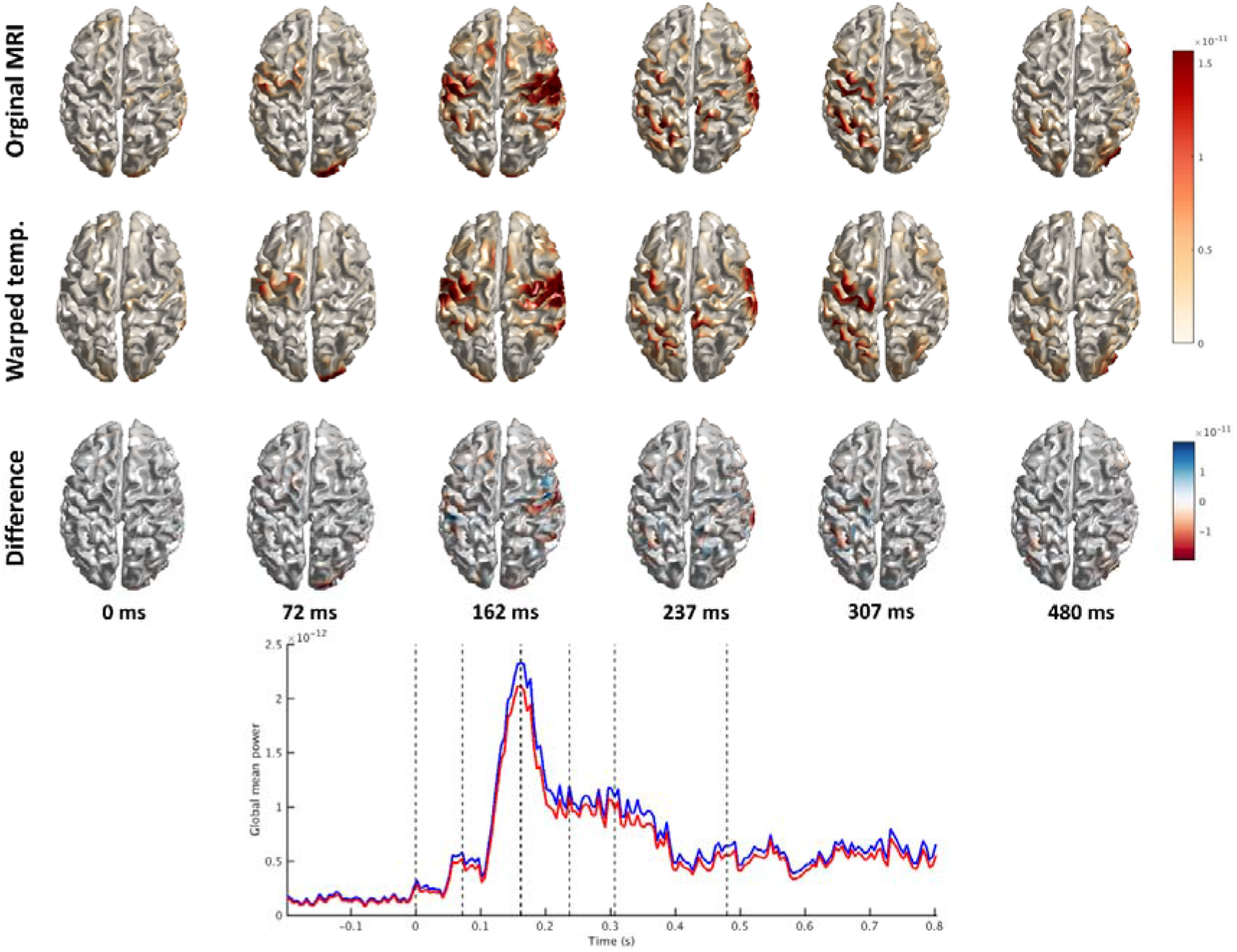
Minimum-norm source reconstruction of the evoked response. Top: Cortical surface topographies of estimated source power for the original MRI (top row), for the warped template (middle row), and the difference between source estimates of the original MRI and warped template (bottom row) at the time-point marked by vertical dashed lines in the global mean power in the plot below. The global mean power is the averaged activity of all source points across time, shown for the original MRI (blue) and warped template (red).

### 3.5 Anatomical MRI features

A sagittal view of the original MRI, the warped template, and the unmodified Colin27 template is presented in Figure 2. Table 1 shows a summary of the segmentation into grey matter, white matter, and CSF volumes (Ashburner & Friston, 2005) for the original MRI, the warped template, and the unmodified template. The scaling of the template to create the warped template resulted in a slightly reduced total brain-compartment volume compared to the original MRI. The grey matter volume was most similar, with only 1.7% less grey matter in the warped template than the original MRI. The white matter volume hard virtually the same volume with only 0.4% less white matter in the warped template than the original MRI. However, the CSF volume was different between the original MRI and the warped template, with 19.9% less CSF volume in the warped template.

**Table 1:**
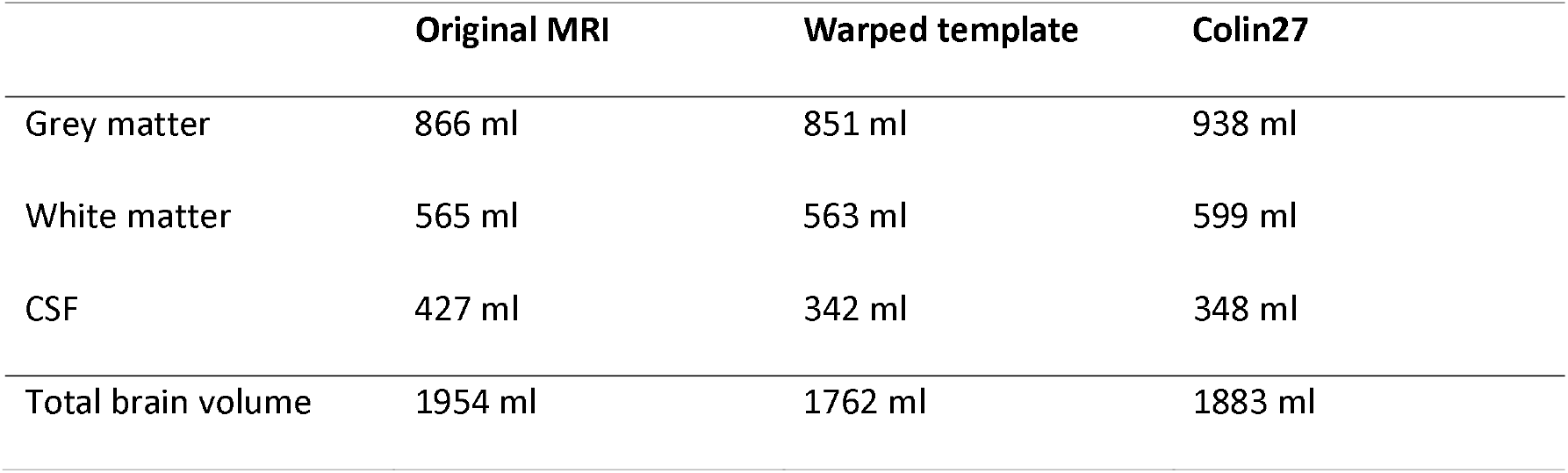
volumes of the segmentation of tissue types: grey matter, white matter, and CSF for the warped template, original MRI and unmodified Colin27 template, and the volume for segmentation of only the “brain” compartment.

|  | Original MRI | Warped template | Colin27 |
| --- | --- | --- | --- |
| Grey matter | 866 ml | 851 ml | 938 ml |
| White matter | 565 ml | 563 ml | 599 ml |
| CSF | 427 ml | 342 ml | 348 ml |
| Total brain volume | 1954 ml | 1762 ml | 1883 ml |

Despite the global volume differences, there was generally a high degree of similarity in the morphological features when compared on a parcel-to-parcel basis. There was high consistency in the surface areas of the parcels between the warped template and the original MRI (α = 0.963) and even excellent similarity in the surface areas between the warped template and the unmodified template (α = 0.991). There was an almost equal consistency in grey matter volume between the warped template and the original MRI (α = 0.967) and between the warped template and the unmodified template (α = 0.987). For the average cortical thickness in the parcellations, there was poor agreement between the warped template and the original MRI (α = 0.633), but still an excellent agreement between the warped template and the unmodified template (α = 0.961), indicating that that the warped template resembles the template rather than the original MRI. The mean curvature within the parcellations showed poor agreement between the warped template and the original MRI (α = 0.434) but still excellent agreement between the warped template and the unmodified template (α = 0.934). Figure 8 shows the summaries of the morphological properties of the original MRI, the warped template, and the unmodified template across all 148 parcellations.

**Figure 8:**
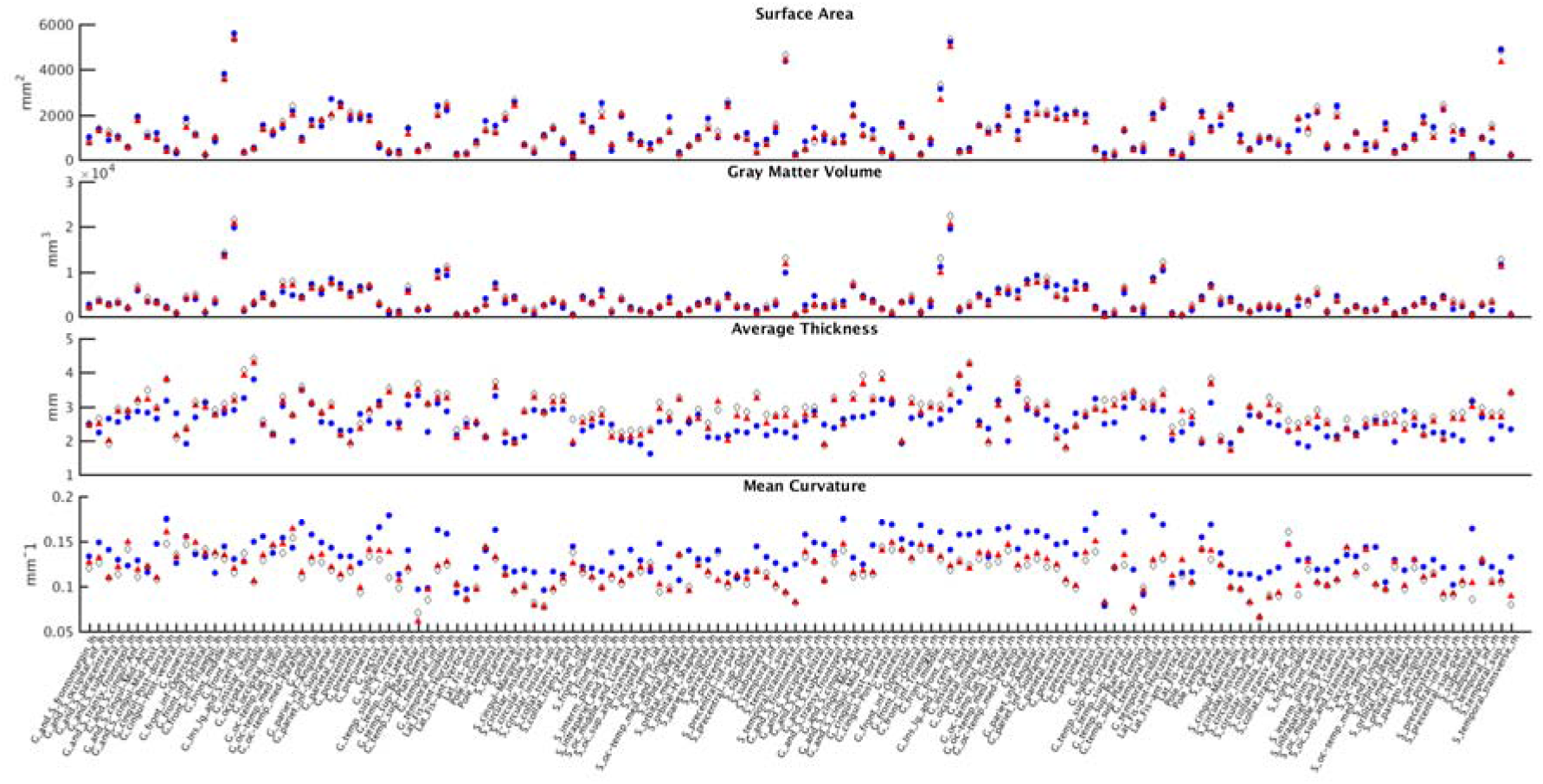
*Surface area, grey matter volume, average cortical thickness, and mean curvature for all 148 parcellations for the warped template (red triangles), the original MRI (blue circles), and the unmodified* Colin27 *template (white diamonds).*

## 4 Discussion

With the example above, we demonstrate that a template MRI warped to the subject’s anatomical MRI can be used to create volume conductor models and source spaces for MEG source reconstruction and still provide results that are quantitatively and qualitatively similar to the results obtained from the analysis using the original MRI. There was a very high similarity between the location and dipole time-series from reconstructed with the warped template and original MRI for the dipole fits. There was a similarly high degree of similarity for the beamformer source reconstructions (DICS and LCMV virtual channels). This was expected since the dipole fits and beamformer methods only incorporate minimal anatomical information. The warped template provide sufficient precision required for replicating previous results and for reusing data in new analyses using dipole fits and beamformer source reconstruction. Therefore, warped templates can serve as an adequate surrogate for data sharing when the original MRI cannot be shared for ethical or legal reasons.

Using the warped template instead of the original MRI for the anatomically constrained MNE source reconstruction resulted in a considerably less similarity between the original MRI and the warped template. This was expected since the MNE method uses the precise anatomical information of the cortical sheet to constrain the inverse solution. Therefore, the MNE results are directly influenced by the geometrical shape of the cortical surface, which differed between the original MRI and the warped template. Therefore, replacing the MRI with warped templates is less suited for this type of whole brain source reconstruction, at least if one aims for high spatial precision. However, for other kinds of analyses, the MNE source reconstructions on the warped templates might sufficiently conserve the general topographical and temporal information compared to the original MRI, allowing certain types of analysis, e.g., analysis focused analysis of activity averaged within an ROI or similar method where millimetre precision of the source reconstruction is not paramount. The use of template cortical sheets is also not uncommon in certain widely used software packages, such as SPM and LORETA; hence we do not believe that this should be disqualified in general.

Whether the degree of similarity of the source reconstructions between source analysis using a warped template or the original MRI is sufficient to replicate a given study or facilitate the reuse of data will depend on the type of analysis one has in mind. For the kinds of analyses requiring light anatomical information (e.g., dipole fits, beamformers), our evaluation indicates the procedure to be adequate for valid (re)analysis, but if the analysis requires more fine-grained anatomical details, the analysis will be less valid. It will be up to the researchers who share the warped template with their MEG data—and the researchers who will use the data for re-analysis—to judge themselves if the shared warped templates are sufficient to carry out the desired analysis.

We do not advocate for the warped template to replace individual anatomical MRIs for constructing volume conductor models and source models when not sharing data, as precision will be lost compared to the original MRIs. For the same reason, any analysis of features in the warped MRIs by themselves, e.g., morphological features such as cortical thickness, folding pattern, etc., is pointless as this is precisely the anatomical information that the pipeline will alter. The proposed pipeline is solely meant for processing MRI for MEG source reconstruction, where it provides a compromise between the need for sharing anatomically plausible data while withholding anatomical details that might breach the confidentiality of the research participants, or that the original researcher is not allowed to share due to lack of consent from the research participants or because of restrictions imposed by the ethics review board.

The use of the warped templates instead of the subject’s real MRIs can be of value when the analysis is constrained by anatomical information but not dependent on the specific morphological features of the anatomical MRIs. Our demonstration focused on conventional SQUID-based MEG source reconstruction, but it is possible that the procedure can be used in sharing other neuroimaging modalities where coregistration between the functional data and structural MRI, e.g., EEG, PET or fNIRS. Though it must be noted that methods which require precise information about tissue outside the brain (e.g., modelling conductivity of the skull and scalp in EEG source reconstruction) will be more affected by dissimilarity between the original MRI and the warped template. In the example presented here, we saw an increase in the bone volume of the warped template compared to the original MRI. MEG signals are, however, virtually unaffected by tissue outside the brain compartment. Conventional SQUID-based MEG sensors are arranged in a fixed arrangement in a cryostat, and the distance between sensors (and consequently the spatial distribution of the magnetic fields due to the cortical activity) depends on the head size. Recent developments in MEG sensor technology allows MEG sensors to be directly placed on the scalp (Andersen et al., 2017; Boto et al., 2018). Consequently, the distribution of the fields depending on the head size might become less in the future. At this moment, the overwhelming majority of MEG systems still uses SQUIDs rather than OPMs and therefore have the sensors in a fixed arrangement in the cryostat.

In the example, we warped the *Colin27* template (Holmes et al., 1998) to demonstrate the pipeline— because it is a readily available template that comes as part of the FieldTrip toolbox (Oostenveld et al., 2011). Any template MRI can, in principle, be used to warp to the subject MRI. It will likely be advantageous to use a template that is suited for the demographics of the participants in the given research project, such as templates that match the age group (Fillmore et al., 2015)—for example, if the dataset includes children, then it will be advantageous use a paediatrics MRI template (Xie et al., 2015)—or use templates matching the ethnicity of the participants (Liang et al., 2016; Rao et al., 2017). It is even possible to use custom templates for the procedure—e.g., in-house templates or an average of all research participants in the study.

The dataset we used to demonstrate the pipeline contained an original MRI that we would characterise as medium quality. In contrast, the template we used is of high image quality averaged over multiple acquisitions (see Holmes et al. 1998 for details). The better quality of the template compared to the participant’s MRI is in this context a disadvantage, as the segmentation process might be biased in favour of the warped template because the template is the same as used in the SPM prior probability maps used for segmentation. The quality of the MRI we used is, in most instances, representative of the quality of the images that researchers have available for MEG source reconstruction. The example, therefore, represents a typical context and data quality where this procedure is of relevance.

Replacing the anatomical information with the template anatomical information should ensure that the research participants cannot be identified based on the MRI data in the shared dataset. It must be noted that sharing the warped templates in place of the original MRI does not fully anonymise the data but only pseudonymised the dataset: as long as there exists a method to link the warped templates to the original MRIs, the data can only be considered pseudonymised. The transformation from template space to individual space links the individual traceable data and the shared warped dataset. It would hypothetically be possible for a third party to identify the research participants if they have access to the transformation—in the same manner, anyone with access to the database that links the participants’ names to the anonymised study code can identify participants. If a third party also has an MRI of the same research participants and would warp the same template to the MRI, they could potentially re-identify the pseudonymised research participants by finding the warped templates with the closest match. However, this hypothetical scenario seems unlikely, and even if so, the warped templates still provide a significant anonymization step compared to sharing only defaced versions of the original MRIs.

While sharing warped templates in place of the original MRIs provides a great deal more anonymity than, e.g., defaced MRIs, we cannot guarantee that the warped template will meet all requirements for data anonymization. Guidelines and regulations for anonymising research data vary between regions and are constantly updated as new methods for anonymization and deanonymization emerge. While we argue that sharing warped templates instead of original MRI provides a great deal more anonymity to the research participants, it is ultimately up to the researchers responsible for data handling and permission to make sure that they have adequate ethical and legal permissions to share data from the relevant institutions and to obtain informed consent by the research participants to share the pseudonymised data. It should also be noted that the pipeline is only about the anatomical images used for MEG source estimation, and the procedure does not solve privacy issues that might arise from other potentially identifying features in the data. It is up to the researchers to consider what data to share and be informed about available anonymization tools and how well these tools remove traceable features.

In conclusion; warping a template anatomical MRI to the subject’s original anatomical can provide similar results in MEG source reconstruction as analysis performed with the original MRI and can therefore provide an adequate replacement for the original MRI when sharing data, as the warped template will not have the same degree of sensitive personal information as the original MRI. The procedure is meant to provide a tool for the seemingly conflicting demand of sharing research data and the demand to keep the privacy of the research participants. We hope that this procedure can help researchers share their MEG data for reproducibility and reuse.

## Authorship contribution statement

**Mikkel C. Vinding:** conceptualisation, methodology, software, validation, formal analysis, investigation, writing - Original Draft, writing - review & editing. **Robert Oostenveld:** conceptualisation, methodology, software, writing - review & editing, supervision.

## Acknowledgements

The authors would like to thank Lau Andersen, Aarhus University for discussions and testing the pipeline.

## References

Aloui, K., Nait-Ali, A., & Naceur, M. S. (2018). Using brain prints as new biometric feature for human recognition. Pattern Recognition Letters, 113, 38–45. https://doi.org/10.1016/j.patrec.2017.10.001

Amico, E., & Goñi, J. (2018). The quest for identifiability in human functional connectomes. Scientific Reports, 8(1), 8254. https://doi.org/10.1038/s41598-018-25089-1

Andersen, L. M., & Lundqvist, D. (2019). Somatosensory responses to nothing: An MEG study of expectations during omission of tactile stimulations. NeuroImage, 184, 78–89. https://doi.org/10.1016/j.neuroimage.2018.09.014

Andersen, L. M., Oostenveld, R., Pfeiffer, C., Ruffieux, S., Jousmäki, V., Hämäläinen, M., Schneiderman, J. F., & Lundqvist, D. (2017). Similarities and differences between on-scalp and conventional in-helmet magnetoencephalography recordings. PLOS ONE, 12(7), e0178602. https://doi.org/10.1371/journal.pone.0178602

Ashburner, J., & Friston, K. J. (1999). Nonlinear spatial normalisation using basis functions. Human Brain Mapping, 7(4), 254–266.

Ashburner, J., & Friston, K. J. (2005). Unified segmentation. NeuroImage, 26(3), 839–851. https://doi.org/10.1016/j.neuroimage.2005.02.018

Ashburner, J., & Friston, K. J. (2009). Computing average shaped tissue probability templates. NeuroImage, 45(2), 333–341. https://doi.org/10.1016/j.neuroimage.2008.12.008

Baillet, S. (2017). Magnetoencephalography for brain electrophysiology and imaging. Nature Neuroscience, 20(3), 327–339. https://doi.org/10.1038/nn.4504

Beckler, D. T., Thumser, Z. C., Schofield, J. S., & Marasco, P. D. (2018). Reliability in evaluator-based tests: Using simulation-constructed models to determine contextually relevant agreement thresholds. BMC Medical Research Methodology, 18(1), 141. https://doi.org/10.1186/s12874-018-0606-7

Boto, E., Holmes, N., Leggett, J., Roberts, G., Shah, V., Meyer, S. S., Muñoz, L. D., Mullinger, K. J., Tierney, T. M., Bestmann, S., Barnes, G. R., Bowtell, R., & Brookes, M. J. (2018). Moving magnetoencephalography towards real-world applications with a wearable system. Nature, 555(7698), 657–661. https://doi.org/10.1038/nature26147

Byrge, L., & Kennedy, D. P. (2019). High-accuracy individual identification using a “thin slice” of the functional connectome. Network Neuroscience, 3(2), 363–383. https://doi.org/10.1162/netn_a_00068

Dale, A. M., Fischl, B., & Sereno, M. I. (1999). Cortical Surface-Based Analysis: I. segmentation and surface reconstruction. NeuroImage, 9(2), 179–194. https://doi.org/10.1006/nimg.1998.0395

Dale, A. M., Liu, A. K., Fischl, B. R., Buckner, R. L., Belliveau, J. W., Lewine, J. D., & Halgren, E. (2000). Dynamic Statistical Parametric Mapping: Combining fMRI and MEG for High-Resolution Imaging of Cortical Activity. Neuron, 26(1), 55–67. https://doi.org/10.1016/S0896-6273(00)81138-1

Destrieux, C., Fischl, B., Dale, A., & Halgren, E. (2010). Automatic parcellation of human cortical gyri and sulci using standard anatomical nomenclature. NeuroImage, 53(1), 1–15. https://doi.org/10.1016/j.neuroimage.2010.06.010

Eke, D., Aasebø, I.E.J., Akintoye, S., Knight, W., Karakasidis, A., Mikulan, E., Ochang, P., Ogoh, G., Oostenveld, R., Pigorini, A., Stahl, B.C., White, T., Zehl, L. (2021). Pseudonymisation of neuroimages and data protection: Increasing access to data while retaining scientific utility Neuroimage: Reports, 1(4), 100053. https://doi.org/10.1016/j.ynirp.2021.100053.

Fillmore, P. T., Phillips-Meek, M. C., & Richards, J. E. (2015). Age-specific MRI brain and head templates for healthy adults from 20 through 89 years of age. Frontiers in Aging Neuroscience, 7. https://doi.org/10.3389/fnagi.2015.00044

Fischl, B., & Dale, A. M. (2000). Measuring the thickness of the human cerebral cortex from magnetic resonance images. Proceedings of the National Academy of Sciences, 97(20), 11050–11055. https://doi.org/10.1073/pnas.200033797

Fox, P. T., & Lancaster, J. L. (2002). Mapping context and content: The BrainMap model. Nature Reviews Neuroscience, 3(4), 319–321. https://doi.org/10.1038/nrn789

Gorgolewski, K. J., Varoquaux, G., Rivera, G., Schwarz, Y., Ghosh, S. S., Maumet, C., Sochat, V. V., Nichols, T. E., Poldrack, R. A., Poline, J.-B., Yarkoni, T., & Margulies, D. S. (2015). NeuroVault.org: A web-based repository for collecting and sharing unthresholded statistical maps of the human brain. Frontiers in Neuroinformatics, 9. https://doi.org/10.3389/fninf.2015.00008

Gramfort, A., Luessi, M., Larson, E., Engemann, D., Strohmeier, D., Brodbeck, C., Goj, R., Jas, M., Brooks, T., Parkkonen, L., & Hämäläinen, M. (2013). MEG and EEG data analysis with MNE-Python. Frontiers in Neuroscience, 7, 267. https://doi.org/10.3389/fnins.2013.00267

Gross, J., Kujala, J., Hämäläinen, M., Timmermann, L., Schnitzler, A., & Salmelin, R. (2001). Dynamic imaging of coherent sources: Studying neural interactions in the human brain. Proceedings of the National Academy of Sciences, 98(2), 694–699. https://doi.org/10.1073/pnas.98.2.694

Hämäläinen, M. S., & Ilmoniemi, R. J. (1994). Interpreting magnetic fields of the brain: Minimum norm estimates. Medical & Biological Engineering & Computing, 32(1), 35–42. https://doi.org/10.1007/BF02512476

Hari, R., & Puce, A. (2017). MEG-EEG primer. Oxford University Press.

Holmes, C. J., Hoge, R., Collins, L., Woods, R., Toga, A. W., & Evans, A. C. (1998). Enhancement of MR Images Using Registration for Signal Averaging: Journal of Computer Assisted Tomography, 22(2), 324–333. https://doi.org/10.1097/00004728-199803000-00032

Koo, T. K., & Li, M. Y. (2016). A Guideline of Selecting and Reporting Intraclass Correlation Coefficients for Reliability Research. Journal of Chiropractic Medicine, 15(2), 155–163. https://doi.org/10.1016/j.jcm.2016.02.012

Krippendorff, K. (1970). Estimating the Reliability, Systematic Error and Random Error of Interval Data. Educational and Psychological Measurement, 30(1), 61–70. https://doi.org/10.1177/001316447003000105

Krippendorff, K. (2018). Content analysis: An introduction to its methodology (Fourth Edition). SAGE.

Larson-Prior, L. J., Oostenveld, R., Della Penna, S., Michalareas, G., Prior, F., Babajani-Feremi, A., Schoffelen, J.-M., Marzetti, L., de Pasquale, F., Di Pompeo, F., Stout, J., Woolrich, M., Luo, Q., Bucholz, R., Fries, P., Pizzella, V., Romani, G. L., Corbetta, M., & Snyder, A. Z. (2013). Adding dynamics to the Human Connectome Project with MEG. NeuroImage, 80, 190–201. https://doi.org/10.1016/j.neuroimage.2013.05.056

Liang, P., Shi, L., Chen, N., Luo, Y., Wang, X., Liu, K., Mok, V. C., Chu, W. C., Wang, D., & Li, K. (2016). Construction of brain atlases based on a multi-center MRI dataset of 2020 Chinese adults. Scientific Reports, 5(1), 18216. https://doi.org/10.1038/srep18216

Nolte, G. (2003). The magnetic lead field theorem in the quasi-static approximation and its use for magnetoencephalography forward calculation in realistic volume conductors. Physics in Medicine and Biology, 48(22), 3637–3652. https://doi.org/10.1088/0031-9155/48/22/002

Oostenveld, R., Fries, P., Maris, E., & Schoffelen, J.-M. (2011). FieldTrip: Open Source Software for Advanced Analysis of MEG, EEG, and Invasive Electrophysiological Data. Computational Intelligence and Neuroscience, 2011, 1–9. https://doi.org/10.1155/2011/156869

Rao, N. P., Jeelani, H., Achalia, R., Achalia, G., Jacob, A., Bharath, R. dawn, Varambally, S., Venkatasubramanian, G., & K. Yalavarthy, P. (2017). Population differences in brain morphology: Need for population specific brain template. Psychiatry Research: Neuroimaging, 265, 1–8. https://doi.org/10.1016/j.pscychresns.2017.03.018

Samuelsson, J. G., Sundaram, P., Khan, S., Sereno, M. I., & Hämäläinen, M. S. (2020). Detectability of cerebellar activity with magnetoencephalography and electroencephalography. Human Brain Mapping, 41(9), 2357–2372. https://doi.org/10.1002/hbm.24951

Schwarz, C. G., Kremers, W. K., Therneau, T. M., Sharp, R. R., Gunter, J. L., Vemuri, P., Arani, A., Spychalla, A. J., Kantarci, K., Knopman, D. S., Petersen, R. C., & Jack, C. R. (2019). Identification of Anonymous MRI Research Participants with Face-Recognition Software. New England Journal of Medicine, 381(17), 1684–1686. https://doi.org/10.1056/NEJMc1908881

Stenroos, M., Hunold, A., & Haueisen, J. (2014). Comparison of three-shell and simplified volume conductor models in magnetoencephalography. NeuroImage, 94, 337–348. https://doi.org/10.1016/j.neuroimage.2014.01.006

Tadel, F. (2021), Using the anatomy templates, accessed 10 November 2021, https://neuroimage.usc.edu/brainstorm/Tutorials/DefaultAnatomy>

Taulu, S., & Simola, J. (2006). Spatiotemporal signal space separation method for rejecting nearby interference in MEG measurements. Physics in Medicine and Biology, 51(7), 1759–1768. https://doi.org/10.1088/0031-9155/51/7/008

Theyers, A. E., Zamyadi, M., O’Reilly, M., Bartha, R., Symons, S., MacQueen, G. M., Hassel, S., Lerch, J. P., Anagnostou, E., Lam, R. W., Frey, B. N., Milev, R., Müller, D. J., Kennedy, S. H., Scott, C. J. M., & Strother, S. C. (2021). Multisite Comparison of MRI Defacing Software Across Multiple Cohorts. Frontiers in Psychiatry, 12, 617997. https://doi.org/10.3389/fpsyt.2021.617997

Tzourio-Mazoyer, N., Landeau, B., Papathanassiou, D., Crivello, F., Etard, O., Delcroix, N., Mazoyer, B., & Joliot, M. (2002). Automated Anatomical Labeling of Activations in SPM Using a Macroscopic Anatomical Parcellation of the MNI MRI Single-Subject Brain. NeuroImage, 15(1), 273–289. https://doi.org/10.1006/nimg.2001.0978

Van Veen, B. D., Van Drongelen, W., Yuchtman, M., & Suzuki, A. (1997). Localisation of brain electrical activity via linearly constrained minimum variance spatial filtering. IEEE Transactions on Biomedical Engineering, 44(9), 867–880. https://doi.org/10.1109/10.623056

White, T., Blok, E., & Calhoun, V. D. (2020). Data sharing and privacy issues in neuroimaging research: Opportunities, obstacles, challenges, and monsters under the bed. Human Brain Mapping, hbm.25120. https://doi.org/10.1002/hbm.25120

World Medical Association. (2013). World Medical Association Declaration of Helsinki: Ethical Principles for Medical Research Involving Human Subjects. JAMA, 310(20), 2191. https://doi.org/10.1001/jama.2013.281053

Xie, W., Richards, J. E., Lei, D., Zhu, H., Lee, K., & Gong, Q. (2015). The construction of MRI brain/hea templates for Chinese children from 7 to 16 years of age. Developmental Cognitive Neuroscience, 15, 94–105. https://doi.org/10.1016/j.dcn.2015.08.008

Xu. W., Kolozsvári, O.B., Oostenveld, R., Leppänen, P.H.T., Hämäläinen, J.A. (2019). Audiovisual Processing of Chinese Characters Elicits Suppression and Congruency Effects in MEG. Front Hum Neurosci. 13(18). https://doi.org/10.3389/fnhum.2019.00018

